# A computational method for direct imputation of cell type-specific expression profiles and cellular compositions from bulk-tissue RNA-Seq in brain disorders

**DOI:** 10.1101/2020.05.28.121483

**Authors:** Abolfazl Doostparast Torshizi, Jubao Duan, Kai Wang

## Abstract

The importance of cell type-specific gene expression in disease-relevant tissues is increasingly recognized in genetic studies of complex diseases. However, the vast majority of gene expression studies are conducted on bulk tissues, necessitating computational approaches to infer biological insights on cell type-specific contribution to diseases. Several computational methods are available for cell type deconvolution (that is, inference of cellular composition) from bulk RNA-Seq data, but cannot impute cell type-specific expression profiles. We hypothesize that with external prior information such as single cell RNA-seq (scRNA-seq) and population-wide expression profiles, it can be a computationally tractable and identifiable to estimate both cellular composition and cell type-specific expression from bulk RNA-Seq data. Here we introduce CellR, which addresses cross-individual gene expression variations by employing genome-wide tissue-wise expression signatures from GTEx to adjust the weights of cell-specific gene markers. It then transforms the deconvolution problem into a linear programming model while taking into account inter/intra cellular correlations, and uses a multi-variate stochastic search algorithm to estimate the expression level of each gene in each cell type. Extensive analyses on several complex diseases such as schizophrenia, Alzheimer’s disease, Huntington’s disease, and type 2 diabetes validated efficiency of CellR, while revealing how specific cell types contribute to different diseases. We conducted numerical simulations on human cerebellum to generate pseudo-bulk RNA-seq data and demonstrated its efficiency in inferring cell-specific expression profiles. Moreover, we inferred cell-specific expression levels from bulk RNA-seq data on schizophrenia and computed differentially expressed genes within certain cell types. Using predicted gene expression profile on excitatory neurons, we were able to reproduce our recently published findings on TCF4 being a master regulator in schizophrenia and showed how this gene and its targets are enriched in excitatory neurons. In summary, CellR compares favorably (both accuracy and stability of inference) against competing approaches on inferring cellular composition from bulk RNA-seq data, but also allows direct imputation of cell type-specific gene expression, opening new doors to re-analyze gene expression data on bulk tissues in complex diseases.

## Introduction

Bulk-tissue RNA sequencing (RNA-seq) yields an average gene expression profile across a collection of heterogeneous cell types, but does not reveal the cell type-specific gene expression profiles within the specific cell populations of interests. Since not all of the cell types are equally involved in disease progression (1), gene expression analysis on the cell types that are most relevant to the disease may be highly desired. For example, developmental processes of organisms including morphogenesis, embryogenesis, and cell differentiation are directly affected by relative composition of cell types (2). Likewise, presence or absence of a particular cell type explains etiology of many diseases (3,4). As an example, Alzheimer’s disease is characterized by changes in the glial populations in the brain (5) while composition of white blood cells can be an indicator of acute cellular rejection of transplanted kidneys (6). It has also been shown how cell type composition plays a critical role in tumorigenesis in which heterogeneity of tumor cells are implicated in cancer metastasis (7). Recent advancement in single cell RNA-seq (scRNA-seq) technologies has made it clear how specific cell types affect the diseases mechanisms. Remarkable findings in autism spectrum disorders (8), schizophrenia (SCZ) (1), age-related macular degeneration (9), and anatomy of human kidneys (10) are a testament to susceptibility of different cell-types in each disease.

Emergence of scRNA-seq technologies has enabled researchers to formalize classification of inherent heterogeneity of cell populations. However, such technologies are more expensive and analytically challenging than bulk RNA-seq assays, confining their employment in population-scale studies. Despite prevalence of experimental approaches to enumerate cells such as laser-capture microdissection and cell sorting, *in-silico* deconvolution is gaining popularity (11). Broadly speaking, computational deconvolution methods can be categorized under two groups including (12) ‘partial’ and ‘complete’ approaches. In the former category, only cellular proportions can be estimated from bulk data while in the latter, cellular proportions and cell-type reference profiles are directly deconvolved from bulk expression data. ‘Complete’ deconvolution approaches can be further split into semi-supervised and unsupervised. Most of the computational methods fall in the semi-supervised category where a set of maker genes for each given cell/tissue types are available (13,14). Another potential classification scheme for *in-silico* cell type deconvolution is based on the type of transcription data whether the method is designed for microarray or RNA-seq data (15). It is unclear whether and how methods exclusively designed for microarray platforms can be effectively adopted for next-generation sequencing data (NGS), given the improved linear associations between true RNA abundance and sequence reads over microarrays (16,17). However, some researchers like Liebner *et al.* (18) emphasize developing RNA-seq-exclusive statistical models.

Given a reference scRNA-seq data from tissues of interest, estimating cellular composition of bulk RNA-seq data as well as estimating cell-specific expression profiles is an important yet challenging computational problem. There have been multiple methods proposed over the past few years such as CIBERSORT (3), CIBERSORTx (19), ABIS (20), MuSiC (21), Deconf (22), lsFit (4), and BSEQ-sc (23). While some of these methods such as Deconf or IsFit can be used in various contexts, others, such as ABIS or CIBERSORT, were primarily developed for certain diseases such as cancers for enumerating immune cell-types and tumor cells. A common feature shared by many of these approaches is their reliance on known markers *a priori* (that is, users need to provide a list of “marker” genes for each cell type) as well as their limited use to specialized and well-studied cell types. Yet, CIBERSORTx provides and additional signature extraction module to generate gene markers to be used during the deconvolution process. In addition, existing methods do not focus on cross-individual genetic variations that may result in different magnitude of variation of “marker” genes in bulk sample from a specific individual. To overcome these limitations and to maximize accuracy of cell-type deconvolution in a data-driven fashion, we introduce CellR (Fig. 1; https://github.com/adoostparast/CellR), a computational method to deconvolve bulk-tissue RNA-Seq data and infer the cellular compositions as well as cell type-specific gene expression values, using an external scRNA-Seq data set as a reference. CellR incorporates cross-individual gene expression variations during the deconvolution process, which assigns different weights to the identified cell markers reflective of variations across individuals in a population. Moreover, given the estimated cellular composition of bulk samples, CellR is capable of imputing expression profiles for each cell type, thus significantly extending the practical utility of the tool beyond cell type deconvolution. Indeed, estimation of cell type-specific gene expression will open new doors to re-analyze gene expression data on bulk tissues in population cohorts on complex diseases, by focusing the comparative analysis on specific cell types. We illustrate a case study how such cell type-specific analysis can generate biological insights beyond traditional bulk tissue-based analysis.

**Figure 1.**
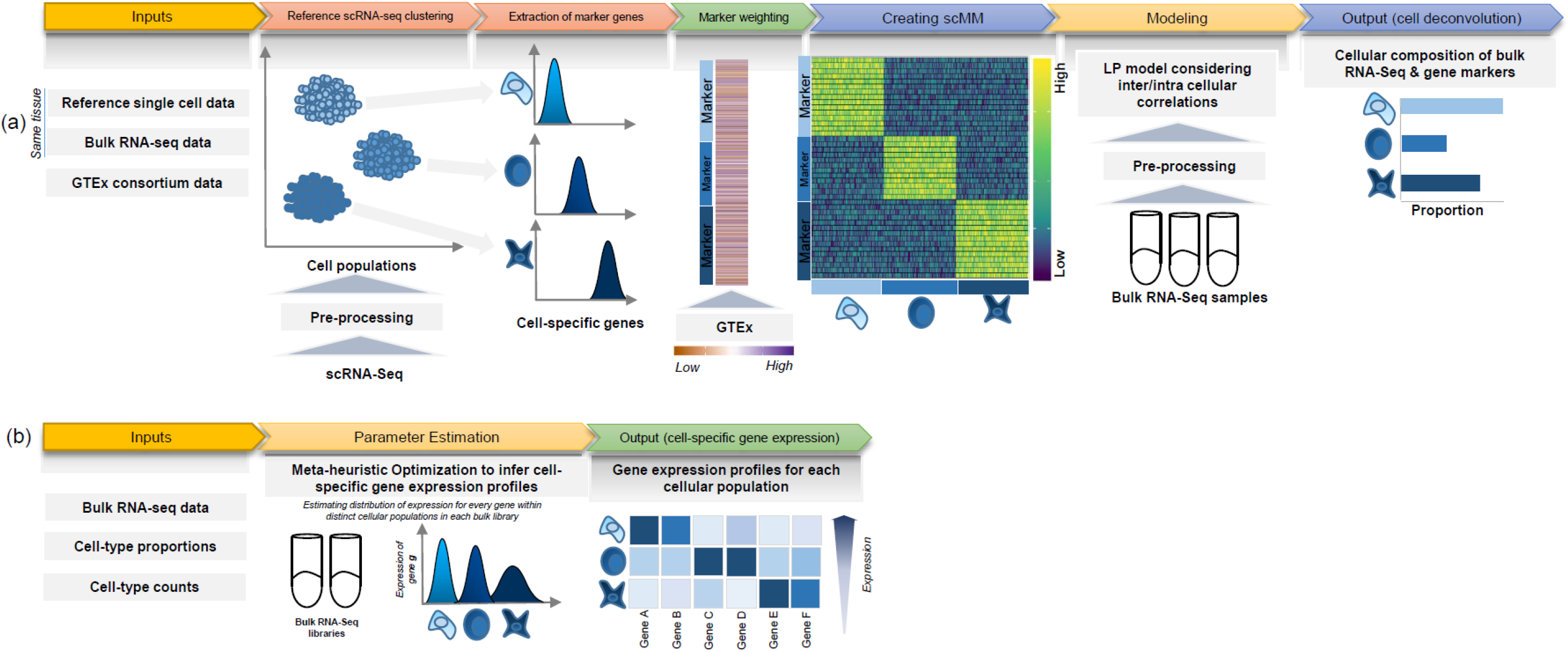
Schematic of the CellR pipeline. (a) Cell deconvolution module: CellR receives the reference scRNA-seq data followed by pre-processing it to remove unwanted artifacts. CellR finds sets of cell types followed by extracting their corresponding markers. In order to account for genetic variations that may modify gene expression, CellR receives TPM matrix from GTEx for the genes from the tissue under study and calculates the weights of the identified gene markers to create scMM. scMM and bulk RNA-seq data, after pre-processing, are fed to the developed linear programming model and cellular composition of each bulk sample will be output; (b) Cell-specific gene expression profiling module: CellR receives bulk RNA-seq libraries, inferred cellular proportions and cell-type counts within each library and processes each library via a newly developed meta-heuristic search optimization algorithm to specify the distribution parameters of each gene within each cell population and outputs a separate transcriptional profiles for distinct cell-types.

## Results

### Model Structure

CellR has two main modules including: (a) cellular enumeration module aimed at estimating the cellular proportions within bulk RNA-seq samples; (b) cell-specific gene expression estimation module which infers the gene expression profiles for each independent cell type of bulk RNA-seq libraries.

In the cellular enumeration module, given the availability of a reference scRNA-seq data from the tissue under study, CellR partitions the cell types and obtains cell-specific genes which are significantly upregulated in each cell type compared to all the others, using Wilcoxon rank sum test. CellR creates a matrix called single cell marker matrix (scMM) describing the expression of the data-derived markers across the sequenced cells while using the cellular annotations provided by the user (see **Methods and Materials**). Next, using the available data from the Genotype-Tissue Expression project, GTEx (24), CellR receives the cross-individual gene expression from specific human tissues and weights the extracted markers so that stable markers, which are less prone to inter-individual variations, rank higher (see **Methods and Materials**). GTEx project is a comprehensive public resource to study tissue-specific gene expression and regulation. Upon applying the obtained weights on the scMM followed by receiving and pre-processing the bulk RNA-seq data to normalize for library size, CellR creates a linear programming (LP) model penalized over the contribution of every single cell in the reference data. Two penalty modes are considered including 1) Lasso mode where contribution of transcription-wise correlated cells, i.e., most of the cells belonging to the same cell type, are shrunk to zero and the most informative cells are used in the model; 2) Ridge mode in which contribution of clustered cells are tightened together so that the overall objective function is minimized (see **Methods and Materials**). After solving the optimization model, cellular proportion of the identified cell types in bulk tissue RNA-Seq data will be given by CellR. Additionally, using the output cellular proportions by CellR, one could generate the predicted gene expression profiles for each cell type, given the bulk tissue RNA-Seq of the sample.

Cell-specific gene expression estimation module receives cellular proportion in a bulk RNA-seq sample, either generated by CellR or similar approaches, consists of a meta-heuristic multivariate search mechanism to optimize the distribution parameters of each gene within each independent cell population which later can be used for downstream analysis (see **Methods and Materials**). This module outputs the overall expression profiles across certain cell populations similar to a bulk RNA-seq data which contains a single cell-type.

### Numerical Experiments on Simulation Data

To test the efficiency of CellR, first, we created two artificial bulk RNA-seq data (see **Methods and Materials**) using two sets of independent scRNA-seq from Lake *et al.* (25) and Segerstolpe *et al.* (26) on cerebellum and pancreas, respectively. We used the procedure recommended by Wang *et al*. (21) to create the artificial bulk data. These two datasets contain 36,166 and 2,209 cells, respectively. The main advantage of such an approach is that the correct proportion of available cell types are already known so that different computational approaches can be evaluated against the known truth. The data on cerebellum contains >5,600 cells including neuronal cell types such as granular cells (Gran, Percentage=58.8%) and Purkinje cells (Purk, Percentage=17.8%) as well as non-neuronal cells including endothelial cells (End, Percentage=1.2%), astrocytes (Ast, Percentage=9.9%), oligodendrocytes (Oli, Percentage=3.4%), pericytes (Per, Percentage=0.77%), oligodendrocyte precursor cells (OPCs, Percentage=5.13%), and microglia (Mic, Percentage=3%). The data from pancreas is a less heterogeneous set of endocrine cells comprising five cells types called *α* (Percentage=60.1%), *β* (Percentage=18.3%), *δ* (Percentage=7.72%), *ε* (Percentage=0.48%), and *γ* (Percentage=13.4%). We ran CellR using two modes (lasso and ridge) and compared its accuracy with a few existing methods including CIBERSORT, CIBERSORTx, MuSiC, Deconf, and lsFit. To test the accuracy, we split the counts of each gene equally to 10 subsets and adopted the cross validation strategy, such that we trained each method using 9 subsets and ran the model on the remaining subset. During each iteration, accuracy was measured using root mean square error (RMSE) and at the end, the average RMSE was reported (Fig. 2a-b). We observed that CellR in lasso mode, CIBERSORT, and CIBERSORTx outperform the other methods while CellR on Ridge mode does not yield the best performance as measured by average RMSE. MuSiC in both cases performs better than CIBERSORT and CIBERSORTx as well as CellR in ridge mode while showing slight increase in RMSE compared to the CellR in lasso mode. IsFit and Deconf underestimated the proportion of abundant cell types such as Gran in cerebellum and *α* cells in pancreas.

**Figure 2.**
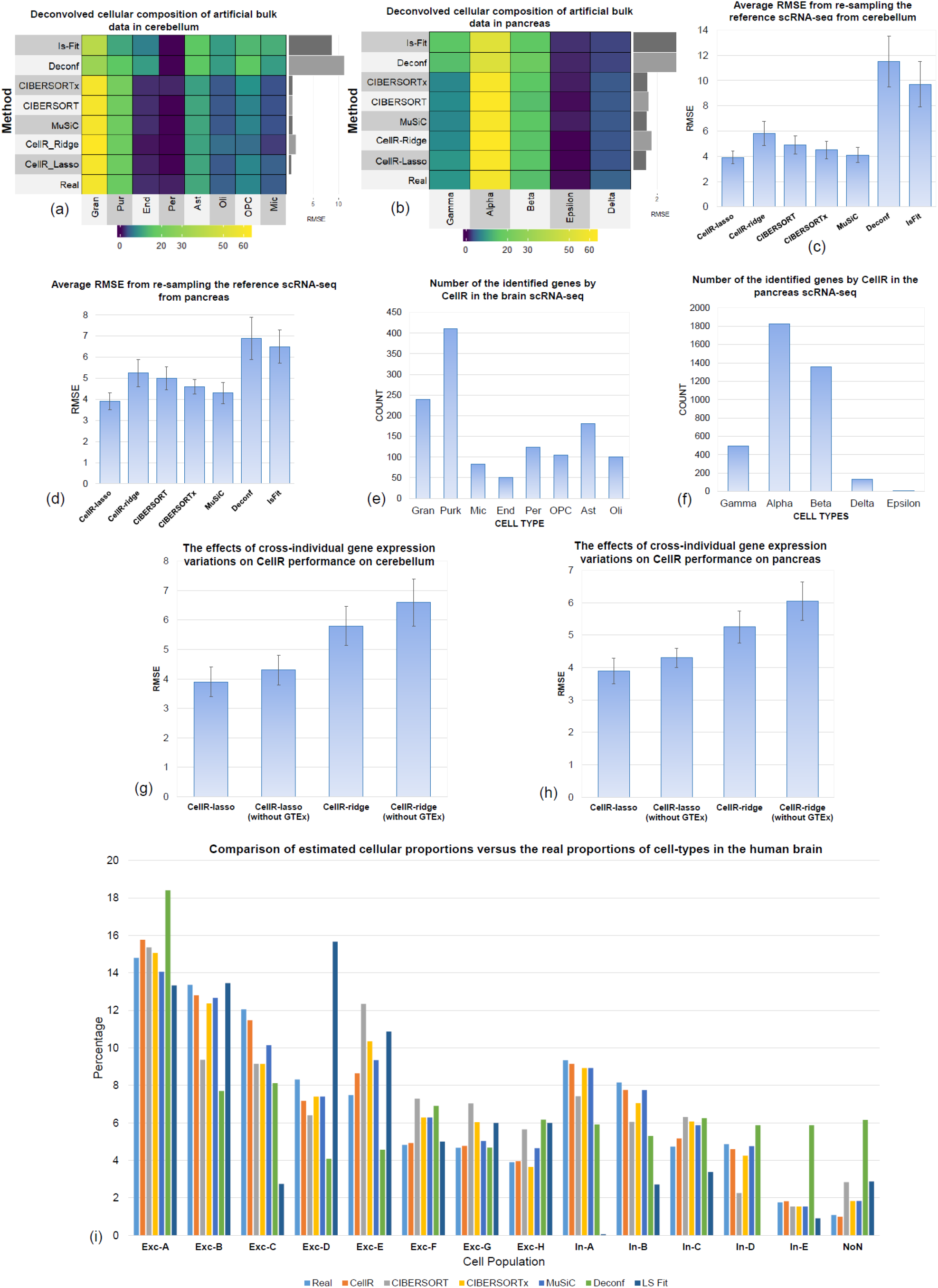
Comparative analysis of CellR and four other competing approaches. (a) Output of the compared methods using the artificial bulk RNA-seq data on cerebellum; (b) Output of the compared methods using the artificial bulk RNA-seq data on pancreas; (c) Average RMSE of re-sampling from the reference scRNA-seq data from cerebellum to compare stability of each method; (d) Average RMSE of re-sampling from the reference scRNA-seq data from pancreas to compare stability of each method; (e-f) Number of the identified cell-specific markers in brain and pancreas, respectively; (g-h) The effects of removing GTEx information from CellR on the accuracy of the results on cerebellum and pancreas data, respectively (i) comparison results of the competing method compared to the ground truth data in human brain.

Additionally, it is known that computational methods for estimating cellular composition may be unstable when the number of cells are small. To compare the stability of the outputs of each method, we re-sampled the reference scRNA-seq data, including 30% of the entire cells in each iteration, performed the experiment 1,000 times and compared the average RMSEs (Figs. 2c-d). CellR in the lasso mode yields more stable numbers with less variation compared to the competing methods. As depicted in Figs. 2c-d, CellR leads to lower RMSEs. The bars representing the average RMSE values for each method includes an error bar. The error bars denote the stability of RMSEs in each iteration which demonstrates CellR to show a reasonable degree of stability compared to the other models.

An advantage of CellR lies in its ability to robustly characterize cellular composition without having a prior biological knowledge of the markers representing cell types. (However, we acknowledge that a prior clustering analysis of the scRNA-Seq data need to be performed to define cellular clusters, which represent cell types). CIBERSORT, on the other hand, requires providing cellular markers that makes it difficult in scenarios where no sufficient information about the underlying molecular signatures of various cell types is known. However, CIBERSORTx provides an automated module to extract gene signatures to be used during the deconvolution process. Details on the identified markers on cerebellum and pancreas are reported in **Supplementary Tables 1-2**, respectively. In cerebellum, CellR revealed 1,292 gene markers while 3,814 genes were identified in the pancreas data. We used the same markers in CIBERSORT. The number of markers per cell-type is provided in Fig. 2e-f.

Another added value of CellR is to consider cross-individual gene expression variations during cell-type deconvolution. To demonstrate this, we repeated the re-sampling procedure described above on cerebellum and pancreas data when cross-individual gene expression variations from GTEx are not used in CellR (Figs. 2g-h). We observed ~9% increase in RMSE for both lasso and ridge modes when GTEx information is not included in the model compared to the cases where GTEx information is available. This stems from uncertainties induced in the linear programming model used by CellR which leads to destabilized outcomes in the optimization stage. Moreover, compared to the other benchmarked methods, it is clear that ignoring the GTEx information worsens the stability of the CellR results and makes a dramatic decrease in the overall accuracy of the method.

An important measure to check the accuracy of the proposed method is to evaluate its performance on sample data where ground truth single cell information is available on the same sample. To this end, we obtained a set of single nucleus RNA-seq data from human cerebral cortex (27) as well as bulk RNA-seq data from the same individual. The data contains 13 cell-types including 8 excitatory (Exc) and 5 inhibitory (In) neurons. We ran CellR and the other competing methods on the bulk data and compared the outcomes with the real number of available cell-types in the bulk library (Fig. 2i). In the majority of the cell-types, CellR yields the most accurate proportions compared to the rest of the methods. In Exc-A set, CIBERSORT and CIBERSORTx perform better while CIBERSORTx and MuSiC show a close performance in the other cell-types compared to the other methods among the competing approaches. Deconf and lsFit demonstrate the poorest performance across the board. Overall, CellR performance was shown to be at a high accuracy in the majority of the profiled cell-types.

### Deconvolution of bulk RNA-Seq data in tissues that are relevant in several diseases

In real experimental situations, reference scRNA-seq and bulk RNA-seq data from the same individual may not always be available. Hence, cell type deconvolution methods should be able to accurately characterize the cellular composition of bulk data coming from different individuals than the source of the scRNA-seq data. To evaluate the performance of our method on real bulk tissue RNA-Seq data sets, using scRNA-Seq data generated on unrelated tissue samples, we obtained two sets of bulk data on postmortem human frontal cortex brain tissues. The first set, provided by Allen *et al.* (28), comprises 278 subjects with the following pathological diagnoses: Alzheimer’s disease (AD), N=84; progressive supranuclear palsy (PSP), N=84; pathologic aging (PA), N=30; control, N=80. The second data was obtained from a study by Labadorf *et al.* (29) on Huntington’s disease (HD) generated from human prefrontal cortex, including 20 HD subjects and 49 neuropathologically normal controls. We used the reference scRNA-seq data from reference (25) on human frontal cortex. We ran CellR as well as four other methods on the two aforementioned datasets. Cellular proportions are reported in Fig. 3a-d. CellR in the ridge mode yields correlated proportions while dispersion of proportions in the lasso mode is relatively higher in astrocytes, inhibitory neurons, and OPCs. CIBERSORT overestimates most of the analyzed cell types both in AD and HD samples. For instance, the proportion of astrocytes given by CellR in AD samples is ~8-21% while the proportion is ~0-58% by CIBERSORT. This is also the case for IsFit and Deconf whose output proportions are overestimated in pericytes. For example, both of these methods report a proportion of over 25% while the real proportion of pericytes in the reference data is less than 1%. Upon making comparisons, we observed that CIBERSORTx tends to yield less dispersed cellular proportions compared to CEIBERSORT. In addition, in most cases, the mean cellular proportions by CIBERSORTx are closer to CellR rather than CIBERSORT including endothelial cells, pericytes, and excitatory neurons in AD as well as microglia, excitatory neurons, astrocytes, and oligodendrocytes in HD. To gain a deeper insight into the number of the identified gene markers within the brain and pancreas scRNA-seq datasets, number of cell-specific markers is shown in Fig. 2e-f. We observed that the number of differentially expressed (DE) genes is larger in cell types which predominantly constitute the overall number of sequenced cells. Our tests indicate that on the simulation data, CellR in lasso mode yields better accuracy while outputting slightly higher dispersion in the proportion of cell types on real data (Fig. 3). As a result, for less heterogeneous data, similar to the simulation data here, we recommend using lasso mode, whereas the ridge mode may have some advantages for more complex real data.

**Figure 3.**
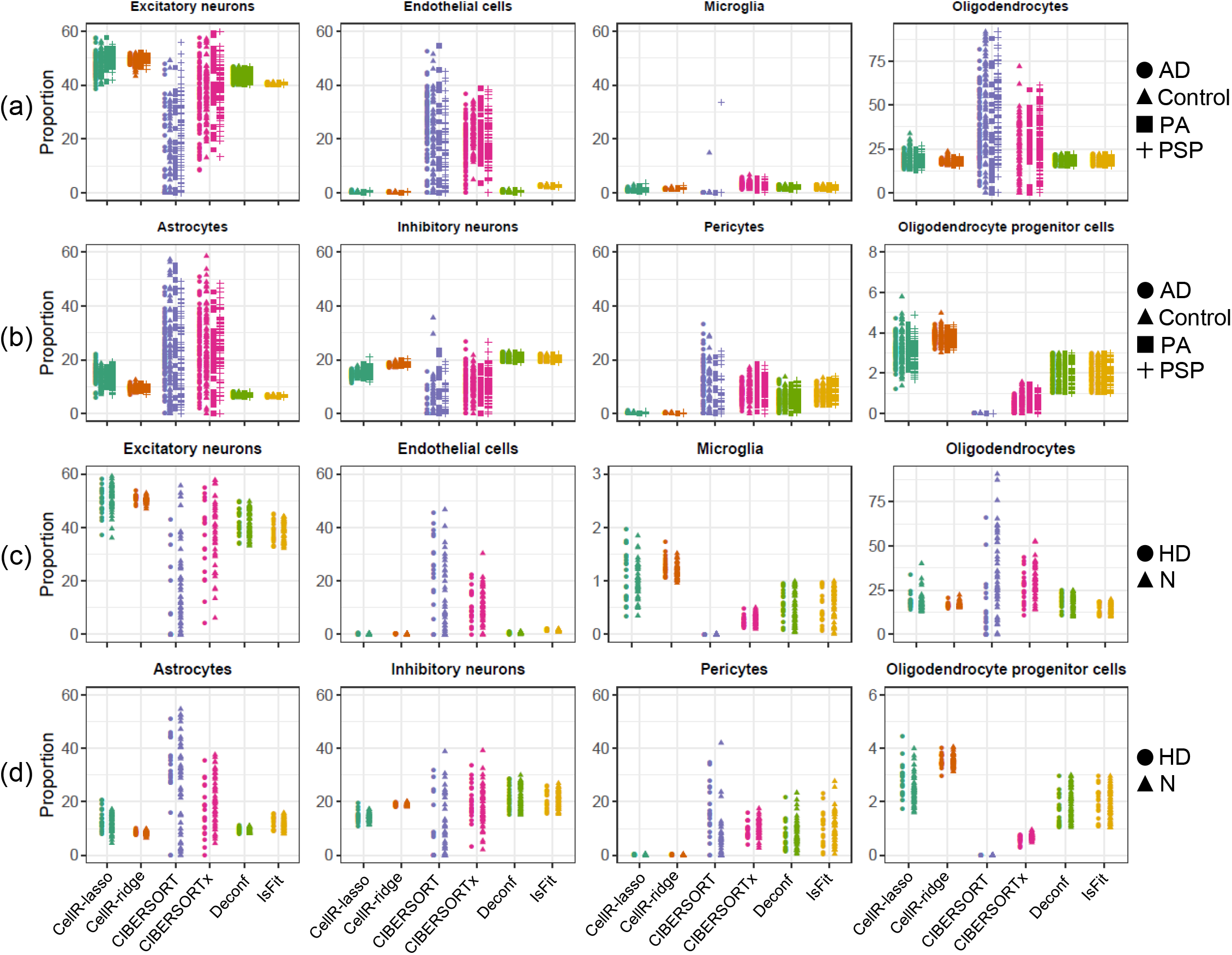
Cellular proportions of AD and HD cohorts; (a) Output of the compared methods using the bulk data from AD samples generated from human brain tissues for Exc, End, Mic, and Oli cells; (b) Output of the compared methods using the bulk data from AD samples generated from human brain tissues for Ast, In, OPC, and Per cells; (c) Output of the compared methods using the bulk data from HD and normal (N) samples generated from human brain tissues for Exc, End, Mic, and Oli cells; (d) Output of the compared methods using the bulk data from HD and normal (N) samples generated from human brain tissues for Ast, In, OPC, and Per cells. N: normal healthy controls; PSP: progressive supranuclear palsy, PA: pathologic aging, Exc: excitatory neurons, In: inhibitory neurons, Ast: astrocytes, OPC: oligodendrocyte progenitor cells, Per: pericytes, End: endothelial cells, Mic: microglia, Oli: oligodendrocytes.

In addition, we analyzed a bulk RNA-Seq data from Fadista *et al.* (30) on type 2 diabetes (T2D), due to the availability of a scRNA-Seq data on pancreas, which is the tissue that is directly relevant to T2D. We used CellR to analyze the putative associations between the proportion of beta cells and HbA1c level, a measure of long-term glycemia. HbA1c denotes normal glucose tolerance (HbA1c ≥ 6.5% in T2D, HbA1c ≤ 6% in healthy individuals). Only CellR successfully captured negative correlations between the beta cell proportion and HbA1c levels (correlation coefficient= −0.41, P-value= 0.003827). We also noticed that a recently published study (21) that re-analyzed the same data has come to a similar conclusion with correlation coefficient = ~-0.31 and P-value=0.00126 (see **Supplementary Figure 1**).

### Estimating cell-specific gene expression profiles

A major application of CellR is to estimate cell type-specific gene expression profiles in distinct cellular populations within a heterogeneous bulk data. We have developed a meta-heuristic optimization-based search mechanism which enables estimating the distribution parameters for distinct cell populations and generate a transcriptomic profile of the cellular constituents of a bulk RNA-seq library (see **Methods and Materials**). CellR receives bulk data, reference scRNA-seq data from the same tissue as well as estimated cellular proportions and counts (whether estimated by CellR or other methods) and generates a starting solution per gene for each cell type. Next, through several layers of search it will estimate near optimal distribution parameters for each gene and generates expression profiles for homogeneous cell populations, separately. To evaluate efficiency of the developed method, we conducted multiple experiments including simulation tests and real-world experiments on SCZ.

Initially, we created 50 pseudo-bulk RNA-seq samples by simulating scRNA-seq data on human cerebellum. For this, we used the data on cerebellum from Lake *et al*. (25) and simulated 50 scRNA-seq datasets (Fig. 4a) upon it using Splatter (31). We turned each simulated data into a pseudo-bulk sample enabling us to have a ground truth for the transcriptome-wide distribution of genes in separate cell-types. Later, we ran CellR on each sample and compared the inferred average expression of the genes with the known expression levels. First, we calculated the similarity of the estimated gene profiles between pairs of cell-types (Fig. 4b) using cosine measure library (see **Methods and Materials**). We had profiled expression levels across 9 cell-types including Gran, End, Ast, Oli, Per, OPC, Mic, as well as two Purkinje cells Purk1 and Purk2. We observed strong similarities between the average expressions of the inferred profiles between the same cell-types (Fig. 4b). For example, inferred profiles of Gran cells indicate a strong similarity with the Gran cells in the ground truth while showing elevated levels of dissimilarity with the other cell-types. Notably, we were able to show how to subpopulations of Purk cells, e.g. Purk1 and Purk2, demonstrate similar patters versus each other while indicating excessive differences with the other cell populations. We also made a second round of comparisons on the basis of our simulations. We calculated the Pearson correlations (Fig. 4c) between the inferred and ground truth expression levels between pairs of cell-types and showed that there is strong correlations between the same cell-types suggesting the reliability of the inferred expression profiles. We acknowledge that the developed method may not be error-free given limitations of scRNA-seq data such as low library depth and dropout effects.

**Figure 4.**
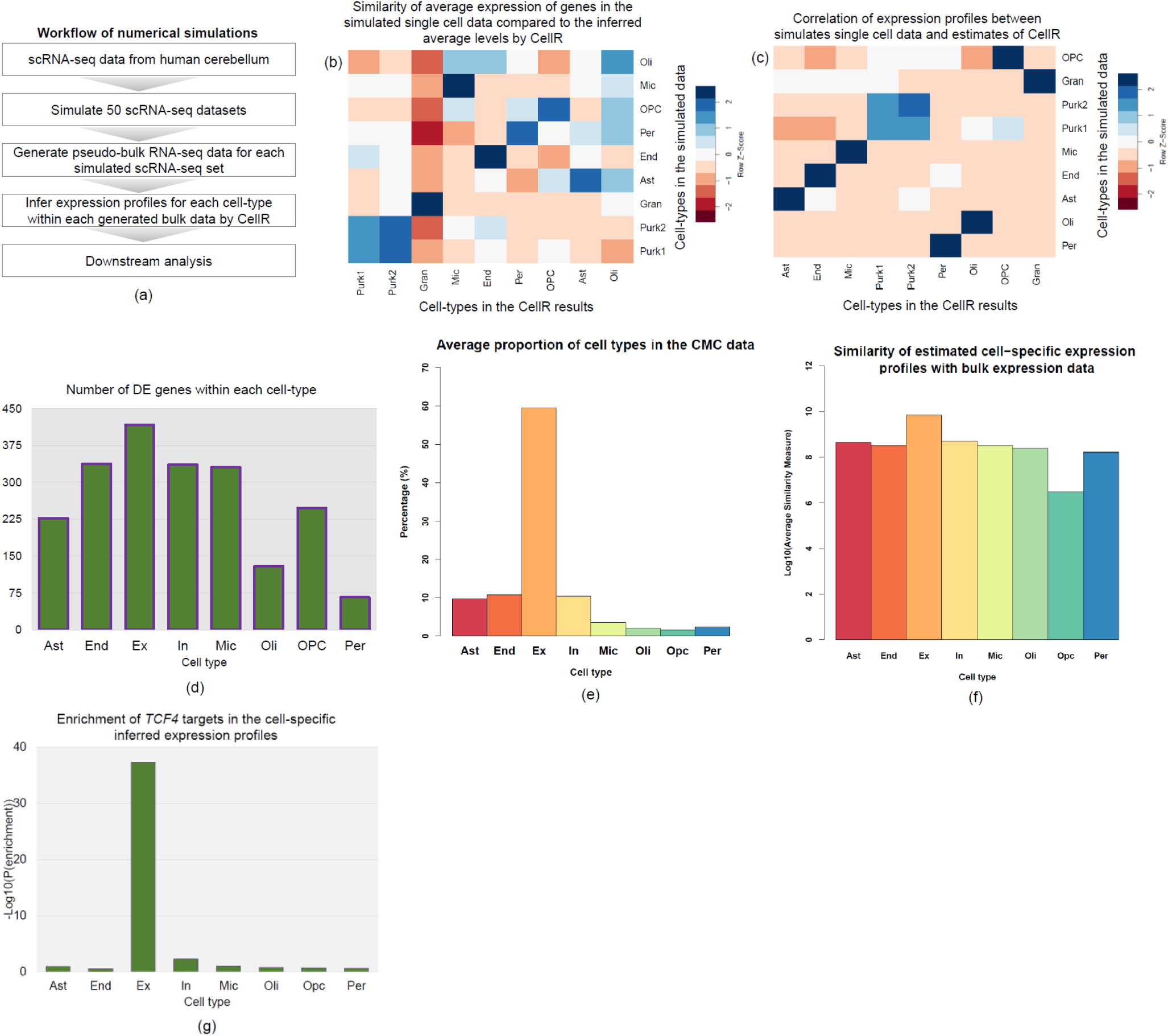
Cell-specific gene expression profiling by CellR. (a) the workflow of simulating RNA-seq libraries to test the efficiency of CellR, (b) similarity heatmap of the inferred gene expression profiles compared to the simulated data on human cerebellum, (c) correlation levels between the inferred gene expression profiles and the simulated data on human cerebellum, (d) number of DE genes within distinct cell populations in the CMC data on SCZ, (e) average cellular proportions in the CMC samples by CellR, (f) average of similarity values of the estimated expression of each gene in a cell-types compared with the bulk data in log10 scale, (g) enrichment degree of the *TCF4* targets being disrupted in the cell-specific expression profiles estimated by CellR..

We were interested to apply CellR on real transcriptome data on SCZ. We used CommonMind Consortium (CMC) study data for this analysis (32). CMC study is currently the largest repertoire of SCZ bulk RNA-seq data on human postmortem dorsolateral prefrontal cortex from a population of 258 SCZ individuals and 279 control subjects. To delineate how transcriptional patterns across distinct cellular populations differ among SCZ and normal individuals, we used CellR to create cell-specific expression profiles on the entire samples in the CMC data on eight cell-types including Ex, Ast, End, In, Mic, Oli, OPC, and Per. We ran CellR and looked for DE genes within each cell-type (**Supplementary Table 3**). Overall, we observed 589 DE genes to be dysregulated in at least one cell-type while 693 genes are DE in the bulk data (Fig. 4d). All of these DE genes were among the DE genes reported in the CMC study. Excitatory neurons found to have the largest number of DE genes (~71% of the total DE genes) while In, End, and Mic cell-types showed an almost identical number of DE genes (~57% of the total DE genes, each). This is consistent with the observations made by Skene *et al*. (1) where Ast and Mic are found to be less relevant in underlying the disease while Ex and In neurons share the highest susceptibility in SCZ.

### Particular cell-types are more relevant to schizophrenia

A study by Skene *et al*. (1) on how common genetic variants in SCZ can be mapped to brain cell types has demonstrated the importance of considering cell-types in studying genetic susceptibility to brain diseases. They had shown that SCZ common variants are predominantly enriched in pyramidal cells, medium spiny neurons (MSNs), and certain interneurons (1). They have concluded that SCZ variants are far less mapped to progenitor, embryonic, and glial cells. A clear picture of susceptibility genes and their corresponding cell-types in SCZ can be achieved by CellR. Therefore, we were interested to evaluate if any of SCZ DE genes can be mapped to certain cell-types. To do so, we obtained the list of 693 DE genes in SCZ from the CMC study (32). Then we used the scRNA-seq reference data by Lake *et al*. (25) and obtained the gene markers by CellR. For each cell-type, we looked for the genes which were shared between their corresponding markers by CellR and the list of DE genes in the CMC data aimed at looking for potential enrichment of DE genes in any of the extracted cell types. We found two cell types of granular cells (P-value=5×10^−3^, fold enrichment=2) and Purkinje cells (P-value=9×10^−3^, fold enrichment=2.4) to enrich for SCZ DE genes. In addition to DE genes, we sought to evaluate whether SCZ common variants are enriched in any of cell-types within the brain. We collected the genome-wide association study (GWAS) hits from the CLOZUK study (33) and the Psychiatric Genomic Consortium study (PGC2) (34) which correspond to 417 protein coding genes that are close to the risk loci (only a fraction of the 417 protein-coding genes may be associated with SCZ though, as GWAS only examine proxy markers of causal variants). We found the same cell-types to be enriched for SCZ GWAS hits including granular cells (P-value=0.022, fold enrichment=2.2) and Purkinje cells (P-value=0.012, fold enrichment=1.8). The rest of the cell types did not pass the significant threshold. Enrichment of SCZ risk factors in certain neuronal cells is in line with the findings of Skene *et al*. (1) where SCZ risk loci were mapped only to neuronal cells. These observations suggest accuracy of CellR in extracting marker genes from reference scRNA-seq data and demonstrates how genetic signals in SCZ originate from neuronal cells. As a proof of concept, we performed extra analysis as follows.

A critical application of CellR is to numerically estimate the proportions of cells-types in bulk samples without conducting costly scRNA-seq experiments. As a proof of concept, using the scRNA-seq reference data on the frontal cortex by Lake *et al*. (25), we obtained the cellular proportions of the samples in the CMC dataset. Average proportions across the entire cohort are represented in Fig. 4e. We clearly see that neuronal cells including excitatory (Ex) and inhibitory (In) neurons, accounts for ~70% of the cellular proportions within each sample. Therefore, we expect that transcriptional signals in these samples predominantly originate from these cell-types. This important observation motivated us to follow how network gene complexes being targeted by SCZ transcriptional master regulators (MRs) are expressed in distinct cell-types. In a recent study (35), we had identified *TCF4* as a SCZ MR through re-analyzing the CMC bulk RNA-seq data and experimentally showed how disrupting expression of this gene can control a large basket of target genes in human induced pluripotent stem cell (hiPSC)-derived neurons. For this, we re-generated cell-specific gene expression levels for each individual in the CMC data (see **Methods and Materials**). To do this, we used CellR to estimate the prior distributions of cell-specific gene expression levels. These cell types include: excitatory neurons (Ex), inhibitory neurons (In), astrocytes (Ast), oligodendrocyte progenitor cells (OPC), pericytes (Per), endothelial cells (End), microglia (Mic), oligodendrocytes (Oli). Then for each distinct cell-type, we generated cell-specific expression levels across the entire individuals in the CMC data, which led to creating eight cell-specific gene expression datasets. Next, for each of these datasets, we created the regulatory networks using the same tools used in our study (35) and obtained the targets of *TCF4*. Finally, we looked for the overlapping targets of *TCF4* generated from the bulk sample versus cell-specific expression data. Only for *TCF4* targets in the data in Ex, we observed a significant overlap (P-val=4.6 ×10^−38^, fold enrichment ratio=119, Fig. 4g). No significant overlap between the *TCF4* targets in the original bulk data versus other cell-specific expression data was observed. We sought to analyze the similarities between cell-specific estimated gene expression profiles of *TCF4* targets and the bulk expression levels. Upon obtaining cell-specific profiles, we calculated the similarities between the estimated expression of each gene in distinct cell-types compared to its expression in the bulk data and averaged the similarity values of the entire *TCF4* targets in various cell-types (Fig. 4f, see **Methods and Materials**). Average similarity of *TCF4* targets in Ex is almost 10 fold higher than other cell-types signifying that the transcriptional signals captured in the bulk RNA-seq data predominantly originates from excitatory neurons. In addition, for each estimated cell-specific expression profiles, we obtained DE genes between SCZ cases versus normal controls and compared them with the list of DE genes in bulk CMC data. We observed significant overlap between the DE genes from Ex-specific expression profiles compared to the bulk data (P-val=2.3 ×10^−38^, fold enrichment ratio=24) while no significant overlap was observed for the rest of the cell-types. All these findings validated the accuracy of CellR in estimating the cellular proportions of bulk RNA-seq data. These observations indicate strong performance of CellR in estimating the cellular proportions and illustrated the importance of taking into account the cellular heterogeneity of bulk RNA-seq data to boost the signals and reduce biological noises.

### Differentially expressed genes are highly enriched in granular and Purkinje cells in Alzheimer’s and Huntington’s diseases

We sought to evaluate if the DE genes in the bulk data can be traced back in the cell-specific molecular signatures, with the hypothesis that DE genes in specific cell types may be the major contributor to overall DE genes identified from bulk RNA-Seq data. To do this, we obtained the list of DE genes between HD samples and negative controls. 5,480 genes have been reported by Labadorf *et al.* (29) to be DE. We intersected the list of DE genes with the identified marker genes by CellR, using scRNA-seq data by Lake *et al.* (25), and found 316 genes by the two groups (Fisher Exact Test (FET) P-Val=0.007). Next, we annotated the shared genes to their corresponding cell-types in the reference scRNA-seq data. 50% of these genes were annotated to Purk and Gran cells which are classified as neuronal cells, whereas the rest of the genes were annotated to five other cell-types. Notably, Purk cells consisted ~33% of the entire set of HD DE genes, e.g. the list of common marker genes by CellR and the DE genes reported by Labadorf *et al.* (29). These cells have been reported to be compromised in aggressive mouse models of HD and their dysfunction is shown to be correlated with HD’s pathology (36). Our observations indicate that a large fraction of DE genes in bulk tissue samples, are in fact markers of specific cell-types. In other words, the statistical signals being picked up in bulk transcriptomic analysis originate from only a fraction of the cellular constituents of the samples further highlighting specificity of cell-types in distinct diseases.

Next, we performed a similar analysis on AD where we obtained DE genes (28) comparing three different pairs including AD-control, PA-control, and PSP-control. No DE genes were observed between PA and normal control samples. We observed 707 marker genes to be DE in AD-control pair while finding 17 marker genes to be DE in the PSP-control pair. We observed ~54% of the DE genes to be enriched in Gran and Purk cell-types, FET P-Val=6.26×10^−32^ . These observations suggest a similar conclusion that much of the signal captured from bulk samples are largely attributed to a limited number of disease-relevant cell-types. Although single-cell sequencing is an effective means to investigate this issue and identify the disease-relevant cell types, it is not cost effective to be scaled to a very large number of samples; in comparison, CellR circumvented this problem and allowed the use of bulk RNA-Seq data to investigate cell type-specific contributions.

## Discussion

Heterogeneous cell populations in many of the genetically-driven diseases are differently predisposed to the disease onset and progression. Such differences cannot be captured by bulk RNA-seq. However, computational deconvolution of bulk mixtures can reveal the proportion of constituent cell-types within the samples. We introduced CellR, a data-driven approach that eliminates the need for having prior biological knowledge on representative gene markers of cell-types (though a prior clustering of the scRNA-Seq is required where each cluster represent a separate cell type) while correcting for potential rare and common genetic variations in the populations that may introduce confounding expression artifacts. As a proof of concept, we made exploratory tests on multiple complex diseases including schizophrenia, Alzheimer’s disease, Huntington’s disease, and type 2 diabetes. We showed how CellR can be effectively employed to yield biological insights into the cellular mechanisms of complex diseases.

Compared to other computational approaches, we demonstrated several unique aspects of CellR in the study: First, CellR outputs more stable proportion values for different samples in the same study. Second, the improved accuracy and lower variation in the identified cell-proportions from CellR demonstrated that we can infer novel biological insights from bulk RNA-Seq samples, as demonstrated in several disease-relevant data sets in our study. Third, with the exception of MuSiC, which considers person-to-person gene expression variations at the single-cell reference level not at bulk resolution, existing methods ignore the variations of gene expressions that differ across individuals, which is a critical factor in elucidating true cell proportions in bulk transcriptomic data. In comparison, the CellR tool makes use of existing knowledge in GTEx knowledge portal to account for cross-individual genetic variations leading to fluctuations in gene expression. In addition, using the identified cellular proportions on SCZ bulk RNA-seq data, we adjusted the gene expression values for distinct cell-types and showed how a significant portion of biological signals in bulk transcriptional signals originate from only excitatory neurons signifying the importance of taking into account the heterogeneity of data when conducting transcriptome studies. Moreover, CellR is designed to estimate near optimal cell-specific gene expression profiles from RNA-seq libraries. Conducting rigorous numerical experiments, we showed how CellR can specify transcriptional dysregulations within distinct cell populations where conventional RNA-seq technologies are not able to distinguish.

We recognize that there are several areas of future improvements that can be incorporated into CellR. First, although our method does not require prior information on specific gene markers, it is possible that well-validated and well-characterized prior information can improve performance. Therefore, we will explore different weighting schemes that allows CellR to take into account the contribution of user-defined gene markers in data analysis. Indeed, some software tools already compiled such a list of gene markers for specific cell types, and we may be able to directly use these as prior knowledge to improve CellR’s performance. Second, as noted by Kong *et al.* (37) that the addition of cell-type proportions as covariates can affect the number of DE genes in bulk data, we envision to take into account such latent knowledge to further reveal the role of cell-specific signals which contribute to the disease progression, *in-silico*.

In conclusion, we developed CellR, a novel computational method to enumerate bulk-tissue RNA-Seq data, infer the cellular compositions and estimate cell type-specific gene expression profiles. Through analysis on simulated data sets and several real data sets on various diseases, our observations corroborate how transcriptional signatures of complex diseases such as schizophrenia, Alzheimer’s disease, and Huntington’s disease are enriched in specific cell-types identified by CellR. Comparative analysis demonstrated better performance of CellR against competing approaches that rely on a few known cell-specific gene markers.

We acknowledge that CellR, given it clustering-based nature, can be influenced by the accuracy of clustering analysis and therefore is not guaranteed to yield the perfect partitioning specifically in highly complex datasets. We expect that CellR can be used to re-analyze many previously published bulk RNA-Seq data and infer more refined biological insights into the cell type-specific contribution of gene expression to disease phenotypes.

## Methods and Materials

CellR is a data-driven method to recover the cellular composition of bulk RNA-seq samples given a scRNA-seq data (usually generated on a different sample but from the same tissue of interest) as a reference. In the following, various stages of CellR depicted in Fig. 1 are thoroughly discussed.

### Optimization model

Let ***f*** be the objective function of the proposed model as follows:

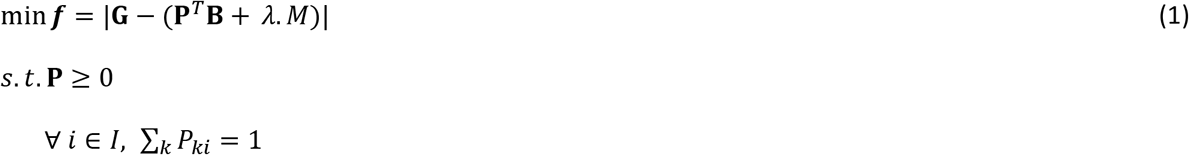

where **G** = []_*I*×*T*_ represents the gene expression levels of the total number of bulk samples (*I*) such that *T* denotes the number of marker genes, *g*_*it*_ represents the expression of marker gene *t* in the sample *i*, **P** = []_*C*×*I*_ represents the proportion of the total number of cells (*C*) in the bulk sample, in which *P*_*ki*_ is the proportion of the cell *k* in the bulk sample *i*, **B** represents the created single-cell marker matrix (scMM), *λ* is the complexity factor, and *M* is the elastic net penalty described in what follows. Extending Eq. 1, the penalty term will be as follows:

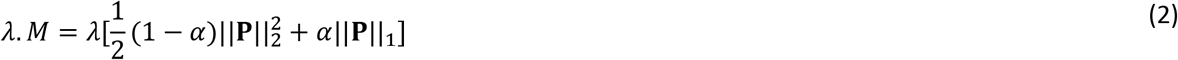

where 0 ≤ *α* ≤ 1. *α* = 0 equates to ridge mode and *α* = 1 denotes lasso mode. In Eq. 4, 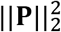 and ||**P**||_1_ denote the *l*_2_ and *l*_1_ norms of the **P** matrix.

In the current version, CellR internally adopts glmnet software package (38) (v. 2.0-16) to solve the optimization problem and uses edgeR (39) (v. 3.22.5) for normalizing the bulk RNA-seq data. glmnet employs cyclical coordinate descent by successively optimizing the objective function over the designed parameters while keeping the others fixed and proceeds the cycle until convergence. Standard procedure recommended by edgeR developers were used to normalize the raw bulk RNA-seq counts. CellR annotates the identified clusters using the cell annotations provided by the user as an input. After solving the optimization model, cellular proportion of the identified cell types in bulk tissue RNA-Seq data will be given by CellR.

### Obtaining expression stability of genes using GTEx data

Let **A** = []_*C*×*T*_ be the matrix of *T* extracted markers from the reference scRNA-seq data across the entire number of cells *C* (CellR internally employs some modules from Seurat (40) for marker extraction). Then, scMM can be obtained as follows:

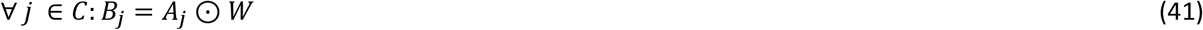

where *W* is the obtained weight vector from Eq. 4, *A*_*j*_ is the *j*-th row of the matrix **A** belonging to cell *j*, and ⊙ represents the dot product of the two vectors. Row-wise concatenation all of the *B*_*j*_ vectors will create the scMM **B**. In order to obtain the weight vector denoted in Eq. 3, let **X** = []_*G*×*In*_ be the TPM (Transcripts Per Million) matrix from GTEx database where *G* denotes the genes and *In* denotes the individuals in the GTEx data. *x*_*ij*_ denotes the expression of gene *i* for the individual *j* in the consortium. Let *X*_*i*_ be the expression of gene *i* across the entire individuals in the GTEx data. We obtain the gene weight vector *W* as follows:

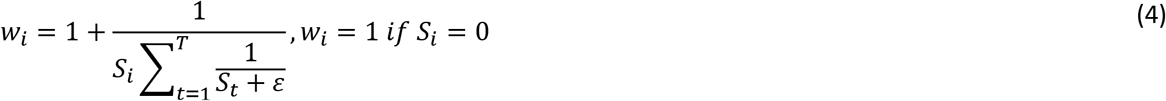

where *w*_*i*_ denotes the weight of the gene *i*, *S*_*i*_ denotes the standard deviation of the expression of the gene *i* across the entire individuals in the GTEx data, *ε* is a very small positive real number to avoid having a zero in the denominator, and *T* denotes the total number of marker genes in scMM.

### Creating artificial bulk RNA-seq data

Suppose **S** = []_*G*×*C*_ be a scRNA-seq matrix containing *C* cells and *G* genes, respectively. The artificial bulk data *B* = []_*G*×1_ can be obtained by summing up the raw counts of each gene across the entire cell population:

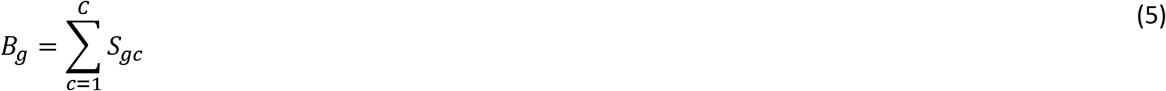

### Competing methods

Three methods were used for comparing the efficiency and accuracy of CellR including CIBERSORT(3) v. 1.06 (https://cibersort.stanford.edu/), Deconf (22), and IsFit (4). We used CellMix 1.6 software package (12) (http://web.cbio.uct.ac.za/~renaud/CRAN/web/CellMix/) in R to run Deconf and IsFit methods. For all of these methods, standard running procedures were applied. To create signature marker list in CIBERSORTx, we used the reference scRNA-seq data as well as the phenotype class files and followed the CIBERSORTx standard procedure to create the signature gene file. We used Seurat (40) to extract the marker genes for the identified cell clusters and used these genes as the input markers for Deconf and IsFit.

### Datasets and preprocessing of scRNA-seq data

The scRNA-seq data on cerebellum was generated by Lake *et al.* (25) and was downloaded from gene expression omnibus (GEO) under accession number GSE97942. The scRNA-seq data on pancreas by Segerstolpe *et al*. (26) was downloaded from ArrayExpress (EBI, https://www.ebi.ac.uk/arrayexpress/) under accession number E-MTAB-5061. The bulk RNA-seq data on HD was generated by Labadorf *et al.* (29) which is available in GEO under accession number GSE64810. The Bulk RNA-seq data on AD and its associated neurological diseases were obtained from AMP-AD knowledge portal (https://www.synapse.org/#!Synapse:syn2580853/wiki/409840) using Synapse ID: syn3163039. To pre-process the raw scRNA-seq count data, CellR internally employs Seurat software (40) (v. 2.3) in R. During the pre-processing stage first, the percentage of mitochondrial gene counts is detected. Then, to normalize the gene expression measurements for each cell, global-scaling normalization is applied followed by multiplying the counts by a scale factor of 10,000 as well as log-transforming the results. Next, the data is scaled by regressing out the percentage of mitochondrial gene content. Using the pre-processed data, principal component analysis (PCA) is done. The number of principal components to be used in clustering for finding the cluster markers can be determined by a resampling test inspired by the jackStraw procedure (42).

### Cross-validation and re-sampling strategies

10-fold cross validation strategy was used to compare the accuracy of CellR against the other methods. First, we split the created artificial bulk RNA-seq datasets into 10 different subsets. In each iteration of cross validation, CellR was applied to each subset and the corresponding RMSE was calculated. Next, the obtained RMSE values in each iteration were averaged and reported on each artificial bulk data. To compare stability of the output of the competing methods, we employed a uniform distribution re-sampling with replacement comprising 30% of the cells in the reference scRNA-seq datasets and trained the models. We iterated the re-sampling for 1,000 times and reported the average RMSEs and their variance in the paper.

### Estimating cell type-specific gene expression

We generated cell type-specific gene expression levels through estimating the prior distributions of each cell-type within the bulk samples in the bulk data. We used the following equation to models the expression of a gene in an individual: 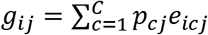 where *g*_*ij*_ denotes the bulk expression of gene *i* in individual *j*, *p*_*cj*_ denotes the proportion of cell type *c* in individual *j*, *e*_*icj*_ represents the expression of gene *i* in cell type *c* for individual *j*, and *C* denotes the total number of cell types. Suppose *e*_*icj*_ to follow a negative binomial distribution of the form *NB*(*r*_*ic*_, *d*_*ic*_). We estimate the parameters of the distributions for every gene *i* across every cell type *c* though a simulated annealing optimization (SA) process (please see the pseudocode in the following). We used the SA structure in our other study (43). The algorithm starts with random initial parameters for expression of each gene regarding distinct cell types. To reduce the risk of falling in local optima, we used the mean of the bulk counts for each gene as the mean parameter of the prior distribution and a randomly generated number between 0 and 1 as the dispersion parameter. During each iteration of the algorithm, using the estimated parameters, sample cell-specific expression values are generated followed by obtaining an estimation of bulk expression levels as follows: 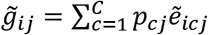, in which 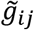 denotes the estimated bulk expression of gene *i* in individual *j* and 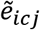 represents the estimated expression of gene *i* in cell type *c* for individual *j* which has been sampled from the simulated prior distribution. Then, in each iteration, root mean square error (*RMSE*_*i*_) is calculated for each gene *i* as follows: 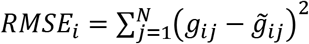 where *N* represents the total number of individual samples in the bulk data. The convergent set of parameters with the lowest root mean square error will then be kept as the prior parameters of cell-specific gene expression levels in the bulk data. We should note that during each iteration of the algorithm for each gene *i*, we have developed two perturbation mechanisms to generate new parameters where each one is randomly selected including: (a) current parameters ± rand(−0.5, .5) × current parameters; (b) new mean parameter (*r*_*new*_): current mean parameters (*r*_*current*_) ± standard deviation of the gene *i* across the bulk data, new dispersion parameter (*d*_*new*_): rand[0, 1]. Another major feature incorporated in this algorithm is its capability to escape from local optimum regions by enabling us to accept parameters (with a restricted probability) with a worse *RMSE* at another domain of the search space to ensure scanning the entire search space for potential global optima (see the pseudocode below).

In the following, we have represented the pseudocode of the developed algorithm:

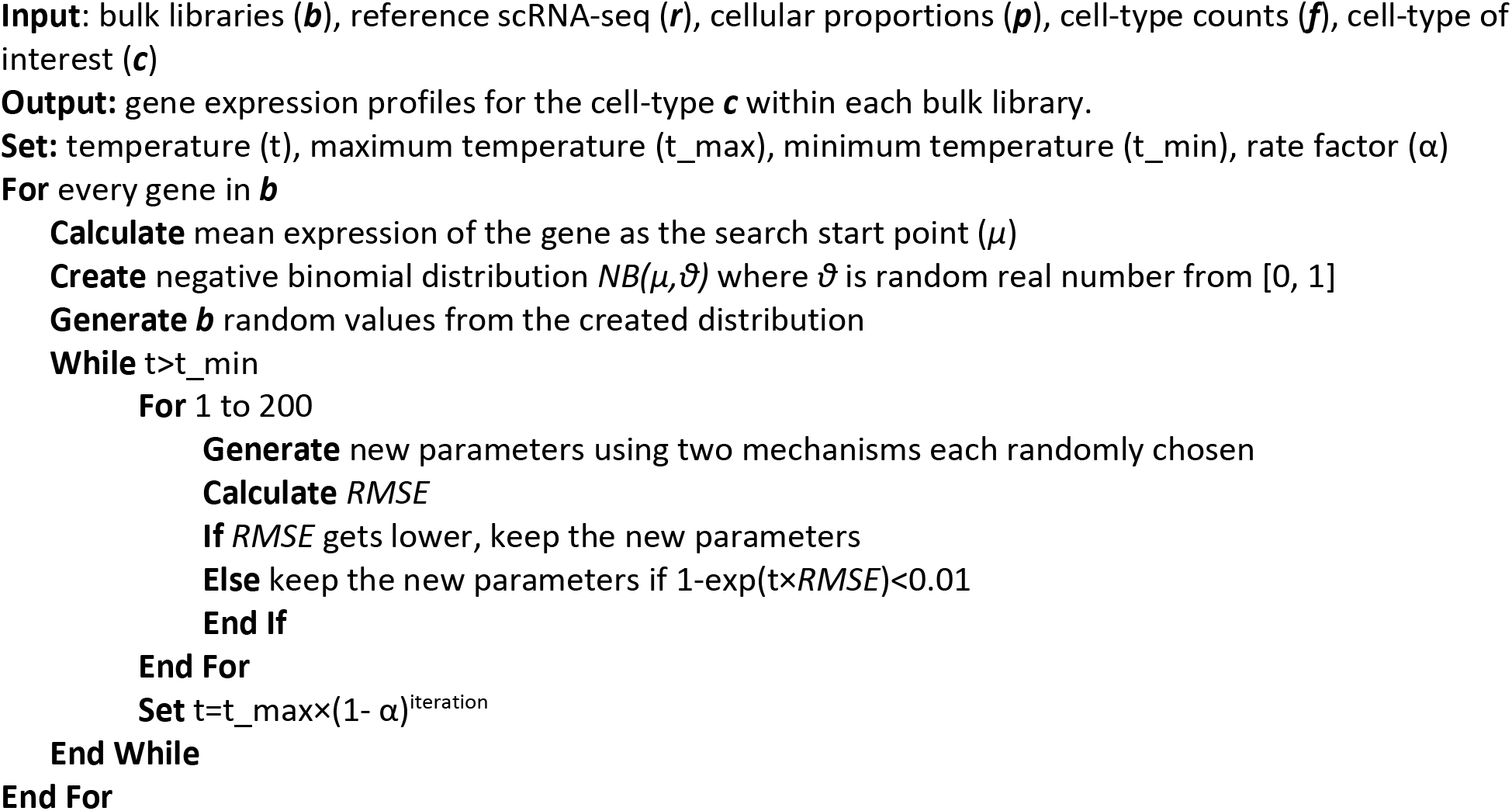

### Measuring similarity of estimated cell-specific expression profiles with the bulk data

In order to measure the similarity of each cell-specific estimation of expression profiles with the original bulk data, we used Cosine similarity measure in text2vec R package. Cosine similarity between two vectors *x* and *y* is defined as follows (44): 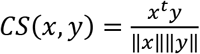 where ‖*x*‖ and ‖*y*‖ denote the Frobenius norm of the two vectors *x* and *y*, respectively. We calculated the similarity of expression profiles of each gene in cell-specific estimated data versus its expression in the bulk data and averaged the similarity values for all of the genes in distinct cell-types.

## Acknowledgement

The RNA-seq data on AD and other neurological disorders were downloaded from AMP-AD knowledge portal under Synapse ID: syn3163039: “Study data were provided by the following sources: The Mayo Clinic Alzheimer’s Disease Genetic Studies, led by Dr. Nilufer Taner and Dr. Steven G. Younkin, Mayo Clinic, Jacksonville, FL using samples from the Mayo Clinic Study of Aging, the Mayo Clinic Alzheimer’s Disease Research Center, and the Mayo Clinic Brain Bank. Data collection was supported through funding by NIA grants P50 AG016574, R01 AG032990, U01 AG046139, R01 AG018023, U01 AG006576, U01 AG006786, R01 AG025711, R01 AG017216, R01 AG003949, NINDS grant R01 NS080820, CurePSP Foundation, and support from Mayo Foundation. Study data includes samples collected through the Sun Health Research Institute Brain and Body Donation Program of Sun City, Arizona. The Brain and Body Donation Program is supported by the National Institute of Neurological Disorders and Stroke (U24 NS072026 National Brain and Tissue Resource for Parkinson’s Disease and Related Disorders), the National Institute on Aging (P30 AG19610 Arizona Alzheimer’s Disease Core Center), the Arizona Department of Health Services (contract 211002, Arizona Alzheimer’s Research Center), the Arizona Biomedical Research Commission (contracts 4001, 0011, 05-901 and 1001 to the Arizona Parkinson’s Disease Consortium) and the Michael J. Fox Foundation for Parkinson’s Research.” The authors would like to thank Drs. Kun Zhang and Blue B. Lake of the University of California, San Diego for generously sharing the RNA-seq data on human brain.

